# Biases in multivariate neural population codes

**DOI:** 10.1101/113803

**Authors:** Sander W. Keemink, Mark C. W. van Rossum

## Abstract

Throughout the nervous system information is typically coded in activity distributed over large population of neurons with broad tuning curves. In idealized situations where a single, continuous stimulus is encoded in a homogeneous population code, the value of an encoded stimulus can be read out without bias. Here we find that when multiple stimuli are simultaneously coded in the population, biases in the estimates of the stimuli and strong correlations between estimates can emerge. Although bias produced via this novel mechanism can be reduced by competitive coding and disappears in the complete absence of noise, the bias diminishes only slowly as a function of neural noise level. A Gaussian Process framework allows for accurate calculation of the bias and shows that a bimodal estimate distribution underlies the bias. The results have implications for neural coding and behavioral experiments.

---

In many brain areas information is distributed across neurons using population codes, in which many neurons respond collectively to a single stimulus. Given its ubiquity, understanding population coding is believed to be crucial to understand coding of information in the brain. By pooling across neurons, population codes allow for accurate estimation of a stimulus from the population response even when neural noise is present. Numerous studies have quantified, among other issues, the role of the tuning curves (Zhang and Sejnowski, 1999), noise-correlations (Sompolinsky et al., 2002; Moreno-Bote et al., 2014), and heterogeneity (Shamir and Sompolinsky, 2006; Ecker et al., 2011; Shamir, 2014) on the coding accuracy.

However, coding accuracy is not the only performance metric. When the same stimulus is repeatedly estimated from a population response and these estimates are averaged over many trials, a systematic difference between the mean estimated value and its true value might remain; this is called bias. In many idealized cases biases are absent from population coding estimation schemes. First, in the limit of low noise, estimators such as the maximum likelihood decoder can be shown to be unbiased (Kay, 1993). Secondly, the coding problem might have an intrinsic symmetry that abolishes bias, that is, over- and underestimation of the stimulus are equally likely - a typical example being the estimation of the orientation of a visual grating from a homogeneous population. Either condition by itself is sufficient to warrant unbiased estimation. For instance, while the maximum likelihood decoder is sub-optimal for high noise, it remains unbiased for one dimensional direction estimations (Xie, 2002).

Yet, in perception biases are common. To explain these, theoretical studies rely on mechanisms that modulate the neural response without adjusting the decoder to break the homogeneity, such as can occur with adaptation (e.g.Stocker and Simoncelli, 2006; Seriès et al., 2009; Cortes et al., 2012) or with contextual changes in the neural tuning (e.g.Schwartz et al., 2007). In contrast to those studies we show that even in homogeneous population codes biases can occur. We consider the case where multiple variables are simultaneously coded in a population, such as occurs in visual cortical area MT when two overlapping transparent random dot motion patterns are presented. We find that in these situations biases in estimation emerge from the decoder itself, even though the decoder has full knowledge of the coding process. Furthermore, when multiple overlapping stimuli are presented, the number of perceived stimuli can be fewer than the number presented, resembling psycho-physical findings (Treue et al., 2000; Edwards and Greenwood, 2005). We develop a mathematical framework based on Gaussian Processes to calculate and understand these effects and discuss their consequences for neural computation and perceptual biases.

## Results

To examine the emergence of biases we consider a population of neurons described by their firing rates. The average response of each neuron is given by its tuning curve *f*(*s*), where s is a vector of stimulus parameters encoded by the neuron. Gaussian white noise *v_i_* with mean zero and variance *σ*^2^ is added to the response, so that on a given trial the firing rate *r_i_* of neuron *i* is

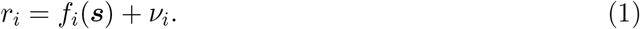

Commonly one studies the case where s is one-dimensional. Here we consider the coding of two stimuli *s* = (*s*_1_,*s*_2_) simultaneously. For concreteness we consider the coding of two overlapping random dot motion patterns in area MT; in this case *s*_1_ and *s_2_* represent the two motion directions, Fig. 1A. The response of MT neurons to such a stimulus has been modeled by the linear average, or, equivalently for our purposes, the sum of the tuning curves to the individual stimuli (van Wezel et al., 1996; Treue et al., 2000),
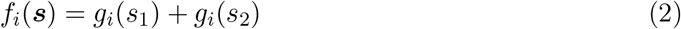

where *g_i_*(*s*) is the bell-shaped tuning of neuron *i* to a single stimulus (Methods). More competitive interactions between the responses have also been proposed; these are considered below.

**Figure 1.**
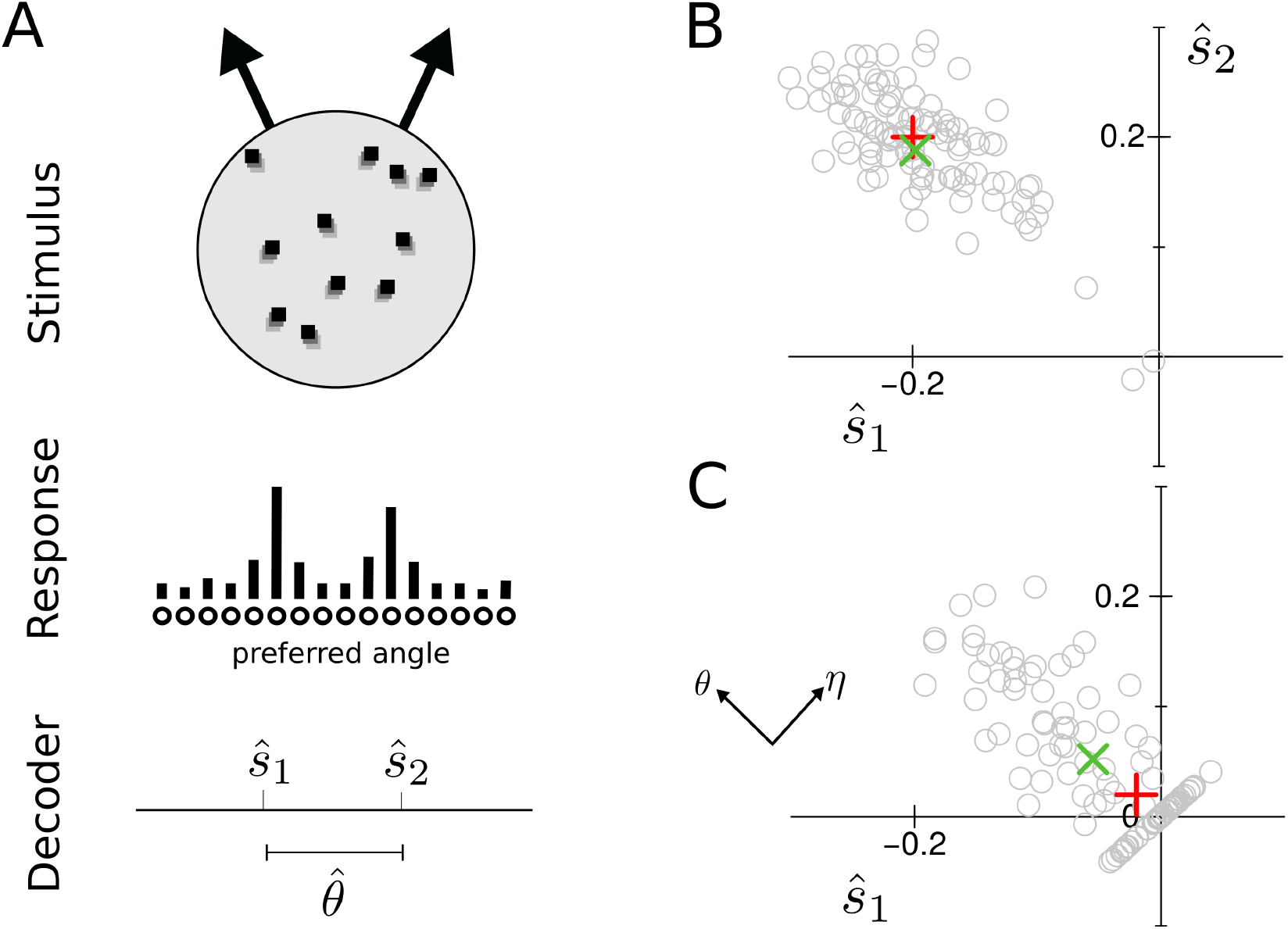
A) Basic encoding-decoding setup. The stimulus consists of two overlapping moving random dot patterns. A population of neurons codes for the two simultaneous stimuli. The task is to estimate the stimulus parameters, here the motion directions *ŝ _1_* and *ŝ*_2_, from the noisy population response. B) Maximum likelihood estimates across a number of trials. For a wide opening angle *s* = (−0.2, 0.2), the distribution of estimates follows approximately a Gaussian distribution. True stimulus (red plus) and average estimate (green X) overlap. C) For narrow opening angles, *s* = (−0.02, 0.02), the distribution of estimates falls into two roughly equal parts, a Gaussian-shaped distribution and a distribution along the line *ŝ* _1_*= ŝ* _2_. True stimulus and average estimate now diverge, i.e. the estimate is biased. The sum and difference angles are indicated by *η* and *θ*, respectively. (all angles in radians).

### Decoding of the neural response

We draw stochastic responses from the above model (see Methods for details) and then decode the stimulus parameters from the noisy population response using the maximum likelihood (ML) decoder. That is, estimates of the stimulus *ŝ*are obtained by finding the stimulus vector that was most likely given the noisy neural response vector *r*,

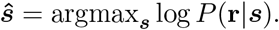

The hat indicates estimates throughout. Because the encoder loses the identity of the stimuli, we additionally impose that *s*_2_ ≥ *s*_1_.

We first consider the case when the two peaks in the tuning curve are far apart (|*s*_1_ − *s*_2_| ≫ *w*, where *w* is the tuning width). In this case the stimulus estimates follow a two-dimensional Gaussian distribution centered around the true stimulus value, Fig. 1B. The true stimulus value (cross) and the mean estimate (X) coincide.

However, when the motion directions are instead almost the same so that the peaks in the population response partly overlap, the distribution radically changes shape, Fig. 1C. Now the estimates fall essentially in two categories: Either the estimates are strongly positively correlated, and cluster on the diagonal where *ŝ*_1_ = *ŝ*_2_. In this case the most likely explanation for the neural response is that the two motion directions are the same. Alternatively, on other trials the estimates are negatively correlated, and the angular difference in the motion direction is over-estimated (repulsion). The mean of neither component of the distribution coincides individually with the true stimulus vector, nor does the mean of the full distribution; in other words, the estimate is biased.

To more easily understand these results we transform the coordinates and describe the system in the sum and difference of the angles. The sum of the angles, *η* = *s*_1_ + *s*_2_ follows a Gaussian distribution and is unbiased as dictated by the rotational invariance of the setup. More interesting, however, is the opening angle Θ = *s*_2_ − *s*_1_. Estimator bias *b* is defined as the difference between mean estimate and true stimulus value, *b*(Θ) = 〈*θ̂*〉 − Θ, where the angular brackets denote the average over trials and *θ̂* are the estimates. The estimator bias is shown as a function of true value Θ in Fig. 2A. When the opening angle Θ is small, the bias is repulsive (the apparent angle is larger than the true value). As the opening angle increases, the bias changes sign and becomes attractive, before reducing to zero for even larger angles, Fig. 2A.

**Figure 2.**
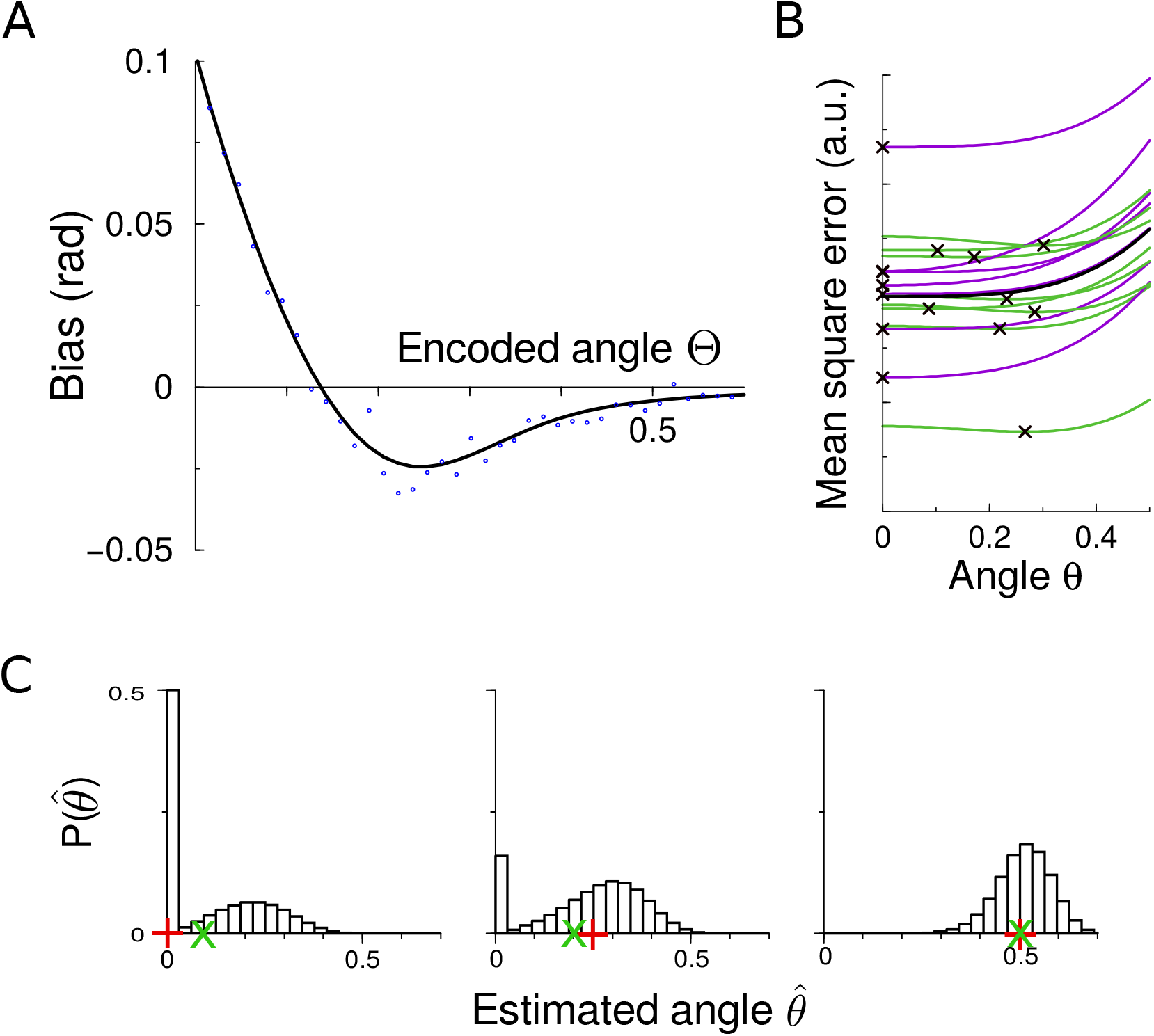
Decoding biases of the opening angle and the underlying decoding distribution. a) Bias in estimation of the opening angle as a function of its true value, showing both a repulsive bias at small angles and a attractive one at larger angles. The curve was calculated using the algorithm given in the Methods. Also shown for comparison are simulations (dots) averaged over 1000 simulations per point. b) Samples of the Mean Square Error in case the true opening angle is zero, the minima of the MSE correspond to the estimate of maximum likelihood estimator. While the average MSE has a minimum at the true value (black curve), on a given noisy trial the estimate can either be exactly *θ* = 0 (shown in purple), or repulsed away from it (shown in green). The black crosses indicate the estimates, i.e. the angle that minimizes the error, on the individual trials. c) Distribution of estimates that underlies the bias. When the true stimulus value is Θ = 0 where the bias is repulsive (left), when the true stimulus value is Θ = 0.25 where the bias is attractive (middle), and when Θ = 0.5 where the bias is virtually absent (right). The true stimulus value is indicated with the red plus on the x-axis, the mean estimate is denoted with the green X. (all angles in radians).

One can wonder whether the repulsive bias is simply caused by imposing *s*_2_ ≥ *s*_1_. However, the distribution of estimates is unexpectedly bi-modal, with a gap between *θ̂* = 0 and the secondary peak, Fig. 2C. Furthermore, the change in the sign of the bias is unexpected from such an interpretation. If the ordering of *s*_1_ and *s*_2_ were randomly assigned, the estimate distribution would become tri-modal with some estimates lying on the diagonal, and others clustering in clouds on the anti-diagonal on either side of the origin.

The bias not unique to the use of maximum likelihood decoder and with a Bayesian decoder similar biases emerge. The Bayesian decoder calculates the full distribution of possible stimulus estimates given the response and the noise model, *P_B_*(*θ*|*r*). For a flat prior for Θ, this is proportional to *P*(*r*|*θ*). Whereas the maximum likelihood decoder takes the maximum of this distribution, using a square loss function the Bayesian estimate equals the mean of this distribution, *θ̂_B_* = ∫ *θP_B_*(*θ*|*r*) *dθ* (Kay, 1993; Salinas and Abbott, 1994). The bias is slightly more pronounced with a Bayesian decoder, Fig S1. However, as the Bayesian decoder does not allow for theoretical treatment, we concentrate on the maximum likelihood decoder.

In summary, in this relatively simple coding problem biphasic biases emerge. Next, we attempt to understand why this occurs.

### Emergence of bias

We now analyze the Maximum Likelihood estimator in detail. For independent Gaussian noise the maximum likelihood estimate is equivalent to minimizing the Mean Squared Error (MSE) E between observed and expected response

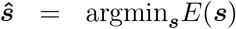

where 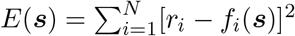. The emergence of the bias and the underlying distribution of estimates can be understood from the mean square error that the estimator seeks to minimize. The MSE is a smooth function but its precise shape varies from trial to trial, Fig. 2B. To write the MSE as a Gaussian Process (Williams and Rasmussen, 2006) we first split it up as

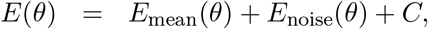

where *C* a stimulus independent term, and *θ* denotes the candidate stimulus. The stimulus dependent part consists of two terms: the first term is the mean 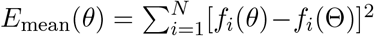 that is identical across trials and attains its minimal value of zero at the true stimulus value, Θ. The second term is the noise term 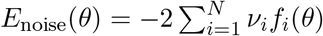.

Of particular interest is the limiting case of Θ = 0. While somewhat contrived as the presented motion directions are identical in that case, exact results can be obtained in this limit that approximately hold for any small Θ. In this limit the term *E*_mean_(*θ*) is lowest at *θ* = 0, as expected, Fig. 2A, black curve. Because of symmetry in the combined tuning curves, Eq. 6, not only all odd derivatives, but also the second derivative of *E*_mean_ is zero. Thus *E*_mean_(*θ*) ~ *O*(*θ*^4^).

The noise term *E*_noise_ is also symmetric and smooth in *θ*, however its second derivative is non-zero. In leading order it is, depending on the noise, either an upward or downward curved parabola centered around the origin. For small *θ* this parabola will dominate over *E_mean_.* Therefore, if the parabola is U-shaped and thus with a minimum at *θ* = 0, the total MSE also has a global minimum there, Fig. 2B, purple curves. If, on the other hand, the noise term has a maximum at *θ* = 0, the global minimum will repulsed away from the true solution, Fig. 2B, green curves. As a result the distribution shows a sharp peak at 0, and a smeared peak further away, Fig. 2C (left). Furthermore, when the encoded angle Θ = 0, exactly half of the estimates will be at *θ* = 0 (i.e. fall on the diagonal in Fig. 1C) and the other half not. As Θ increases, the probability to find estimates *θ̂* = 0 will decrease and the second distribution will gain more mass until the mass at zero disappears, Fig. 2C (middle and right). The net effect is that this will first decrease the repulsive bias, then turn into an attractive bias, and finally the bias disappears.

In the Methods we describe how the Gaussian Process approach can be used to calculate the probability of estimates *P*(*θ̂*|Θ) in a numerically exact way without relying on simulations. This method was used to create Fig. 2A+C, and compares well to explicit simulations over many trials (dots in Fig. 2A).

### Dependence of bias on noise

The bias curve depends on the neural noise level, Fig. 2A and other system parameters. In the limit of small angles the bias can be found by estimating the expected location of the minima of the Mean Square Error (markers in Fig. 2B). As shown in the Methods this gives for the tuning curves used,

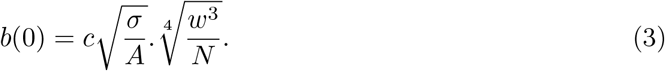

where *σ* is the std.dev. of the neural noise, *w* is the tuning width, *A* is the maximum neural response amplitude, *N* is the number of neurons and *c* ≈ 1.2 is a numerical constant. Therefore to, say, half the bias, one needs 4 times less noise, or 16 times as many neurons. The second effect of the noise level is a shift in the angle at which repulsion becomes attraction, i.e. where the curve in crosses the x-axis (see Fig. 4A). Because the slope of the bias is exactly −1 at the origin (*b*(Θ) ≈ *b*(0) − Θ + *O*(Θ^2^), see below), the location of this transition point is well approximated by the bias at zero.

Interestingly, as the noise is reduced, the distribution of estimates remains bi-modal. While in the limit of zero noise the bias disappears as the theory of maximum likelihood estimation requires, the transition in the limit of small angles is not due to a collapse of *P*(*θ*̂) into a single Gaussian distribution, rather it is due to the two peaks in the distribution of estimates moving closer and closer together.

### Effect of the bias on estimator efficiency

The quality of population code readout is not quantified by the bias alone, but also by the amount of trial-to-trial variations in the estimates, i.e. the variance in the distributions in Fig.1B+C. The variance in the ML decoder estimates follows directly from the distribution of estimates *P*(*θ̂*|Θ) that our Gaussian Process approach yields. The variance in the estimator at a particular noise level is plotted in Fig. 3A.

**Figure 3.**
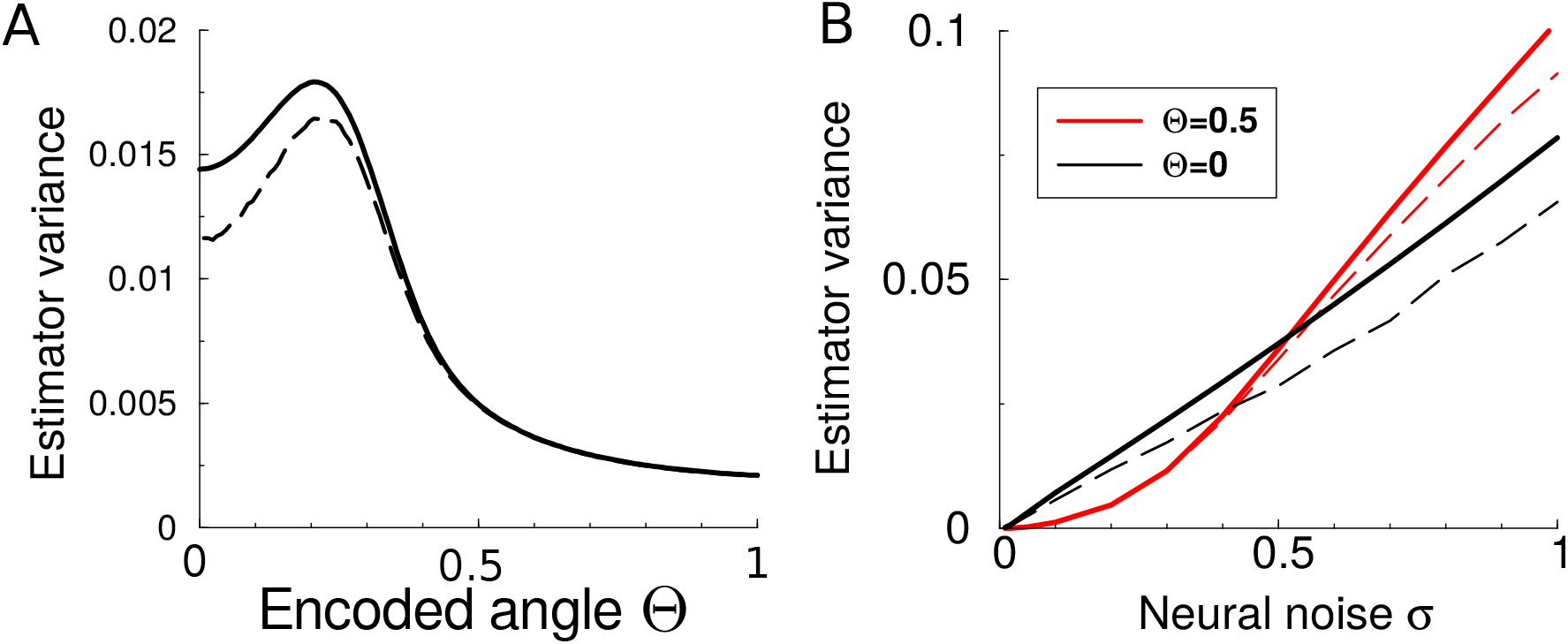
Variance and efficiency of the maximum likelihood decoder and its dependence on encoded angle and neural noise. A. Variance in the estimates depends non-monotonically on the encoded angle. Dashed curve correspond to the Cramer-Rao bound; no estimator can achieve a lower variance. B. Variance in the estimates as a function of the neural noise comparing large and small encoded angles. At small angles the strong bias alters the expected square dependence on noise into linear behaviour. Dashed curves correspond to the Cramer-Rao bound.

The minimal variance any estimator can achieve is limited by the Fisher Information through the Cramer-Rao bound which states that the variance of any estimator obeys (Methods)

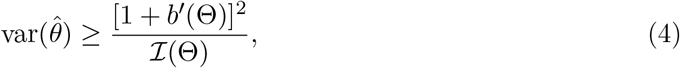

where *b′* is the derivative of the bias, and the Fisher Information 𝓘 (Θ) is given by Eq.8, Methods. The efficiency of an estimator expresses how close it comes to this bound. The resulting Cramer-Rao bound is indicated by the dashed curve in Fig. 3A. The estimator performs at the theoretical limit for large opening angles, and diverges (although it remains close) for smaller opening angles.

For large Θ, the estimate distributions are Gaussian with a width proportional to the neural noise; the bias plays a minor role. As expected from the Fisher Information, the variance of the estimator is proportional to the square of the neural noise, Fig. 3B, red curve. However, near Θ = 0, the bias has a profound effect on the estimator. The estimator variance at Θ = 0 can be approximately found by describing the estimate distribution *P*(*θ̂*|Θ) as a peak at zero and a Gaussian, Fig.2C. As for small angles the spread of the Gaussian is smaller than its mean, the variance is similar to the bias squared, and thus its parameter dependence can be calculated from squaring Eq. 3. Therefore, in contrast to square dependence at large angles, the variance in the estimates is only linear in the neural noise, as is confirmed in Fig.3B, black curve. Finally note that for low noise levels the variance at small angles is larger than the variance at larger angles, but that this switches at high noise.

Interestingly, Cramer-Rao bound follows the estimator performance and is linear in the neural noise, Fig.3B, dashed curves. In contrast, the Fisher Information is proportional to the neural noise squared *σ*^2^, Eq. 8. The reason is that for small angles the Fisher Information goes to zero (Eq.8), but, because the estimator is a smooth, symmetric function, its derivative at the origin equals *b′*(*0*) = −1. Hence at small angles, both numerator and denominator of the Cramer-Rao bound, Eq. 4 go to zero and the net result is a linear dependence on the neural noise. For the parameters used, the MLE always achieves an efficiency ≥ 95%.

As an aside, in calculating the Cramer-Rao bound another advantage of the Gaussian Process approach shows. With simulations the bias and in particular its derivative are hard to calculate accurately, even using a large number of realizations, Fig. 2A dots. However, the numerically exact method to calculate the bias (Methods) allows for a precise calculation of the bias and its derivative. For instance, for a Bayesian decoder the precise bias is much harder to obtain, Fig. S1.

In summary, the bias does not simply lead to a small correction in estimator performance, but fundamentally alters it.

### Competitive coding reduces bias

The estimation bias depends on the encoding model, that is, how the stimuli are coded in the neural response. Above it was assumed that the neural response to two simultaneous stimuli was the sum of the responses to the individual stimuli. While there is some experimental evidence for such a linear interaction, in other studies evidence for more competitive interaction has been found in area MT (Britten and Heuer, 1999), as well as other visual cortices (Gawne and Martin, 2002; Lampl et al., 2004; Oleksiak et al., 2011).

Such interactions have been modeled using a maximum-like interaction, so that instead of Eq. 2, the response of a single neuron to two simultaneous stimuli is

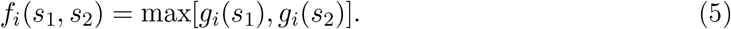

When the simulations are repeated for this encoding model, the bias is still present, but it is substantially smaller, Fig. 3A.ii. The repulsive bias is now approximately linear in the noise and the attractive component of the bias is smaller and becomes only apparent at even higher noise levels (see inset). Also the variance in the estimate is smaller, Fig. 3B, however, the bias still substantially alters the performance limit - compare the Cramer-Rao bound before (dotted) and after bias correction (dashed).

The mathematical reason for the reduced bias and variance, is that in this case the mean term in the MSE is not quartic but quadratic, reducing the bias. Thus we find that the precise coding model is an important determinant of the size of the bias, and these findings suggest a functional role for competitive interactions.

## Discussion

Traditionally, theoretical studies of population codes have focused on estimator variance. Whenever biases have been studied, they have been explained from inhomogeneities in the neural encoding. Here we find that when multiple stimuli are encoded simultaneously in a relatively simple coding problem, substantial biases arise, which unlike previous mechanisms, are intrinsic to the decoder. That biases occur is in itself not surprising. Apart from cases where symmetry rules out biases, the absence of biases can only be proven in the limit of low noise, and in general an ML decoder will not be unbiased (Kay, 1993; Seriès et al., 2009; Pilarski and Pokora, 2015), nor efficient (Xie, 2002). Yet, the rich structure of the biases in these simple models, including its biphasic character and its relative persistence at low noise, is surprising.

The reason for the biases is the bimodal distribution of decoding estimates. By using a Gaussian Process approach we have developed an analytical theory of ML decoders which allows calculation of this distribution and the bias. Although the biases will disappear in the limit of zero noise, the bias diminishes only slowly as noise is reduced (proportional to the square root of the std. dev. of the neural noise). The persistence at low noise contrasts other studies of bias and efficiency where effects disappear abruptly when noise is lowered (Kay, 1993; Xie, 2002).

The results generalize in a number of directions. While we have only shown results for additive Gaussian noise, simulations show that our results extend to Poisson and multiplicative Gaussian noise, as well as correlated noise. Competitive encoding, such as the maximum coding, can reduce the bias, but will not abolish it, Fig. 4. Similarly, while our analysis relies on the maximum likelihood decoder, we find that the results are not unique to using the maximum likelihood estimator and occur with Bayesian decoders as well (Fig. S1). The nature of neural decoding mechanism is currently not clear, although it has been argued that it is straightforward to implement ML decoders neurally (Jazayeri and Movshon, 2006).

**Figure 4.**
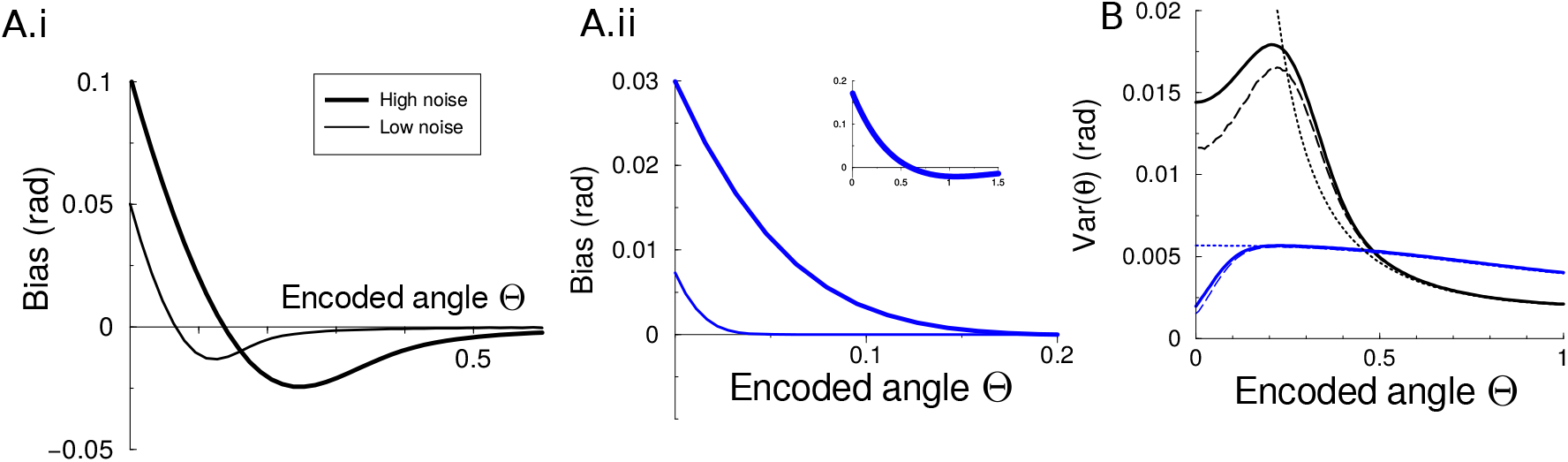
Bias and variance in a competitive coding model. A.i Bias in the estimates in the linear coding model for two different noise levels. The standard deviation of the neural noise was either high (*σ =* 0.2) or low (*σ =* 0.05). A. ii Bias in the estimates in a competitive coding model where the response of any neuron to two stimuli equals the maximum response to the individual stimuli for the same noise levels as used in panel A.i. Only at very high noise levels (*σ* = 1), the attractive bias manifests itself (inset). Note the difference in scales. B. Trial-to-trial variance in the estimates for the linear coding model (black) and the competitive coding model (blue). Dashed curve correspond to the Cramer-Rao bound. Dotted line corresponds to the Cramer-Rao bound uncorrected for the presence of bias (i.e. the inverse Fisher Information).

The results bear upon psycho-physical experiments where two overlapping random dot motion patterns with different directions are presented and subjects are asked to guess the angle between the two directions. In such experiments repulsive biases have commonly been observed (Marshak and Sekuler, 1979, but see Braddick et al., 2002). Several effects have been hypothesized to underlie these biases, including adaptation (the bias *increases* with presentation time,Rauber and Treue, 1999), cortical interactions (Carandini and Ringach, 1997) and repulsion from the cardinal directions (Rauber and Treue, 1998). The bias described here, is not at odds with those explanations, but presents a novel contribution to the total bias that is intrinsic to the decoder and which should be most prominent at small angles and for short presentation times.

The estimate distribution can be seen to reflect an ambiguity between the presence of one or two stimuli. Apart from predicting a bias, the theory predicts a bi-modal distribution of direction difference estimates and for small angles about half the time the two motions should be perceived as one. In experiments the number of stimuli that can simultaneously be perceived using overlapping motions is limited (e.g.Edwards and Greenwood, 2005) and when three or five overlapping motions are presented, they can sometimes be perceived as two (so called metamers,Treue et al., 2000); an effect which previously has been explained using the probabilistic population code framework (Zemel et al., 1998; Zemel and Dayan, 1999). The results here suggest that differences in the numerosity between presented and perceived stimuli already emerge with maximum likelihood decoders. Quantitative verification of this prediction of our study should be possible but might be challenging as participants’ expectations and, similarly, natural priors for perceiving a single motion direction instead of two directions can influence results.

## Acknowledgments

SK was supported by the EuroSpin Erasmus Mundus program and the EPSRC NeuroInformatics DTC. The authors would like to thank Udo Ernst, Richard van Wezel, Lawrence York, Peggy Series, and Chris Williams for discussions.

## Methods

### Neural population response

We use a population of *N* = 100 neurons. The tuning of neuron *i* to a single stimulus is given by 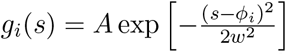. Here *A* is the response amplitude (arbitrarily set to 1), *w* is the width of the tuning curve (set to 1/2). The preferred directions *ϕ_i_* of the neurons are equally spaced between 0 and 2 π. As is common, we assume that the angles involved are relatively small, so that we don’t have to worry their circularity, which would add complication through the need for circular statistics but does not change the results qualitatively.

When multiple stimuli are present, the neural response is modeled as the sum of the responses to the individual stimuli. After transforming the variables to the sum angle *η* and the difference angle Θ (see Main text) we can set *η* to zero, so that the tuning of neuron *i* becomes

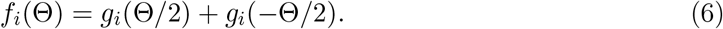

By replacing *A* by half its value one can obtain a model where the joint tuning curve equals the average (instead of the sum) of the tuning curves. The default value of the std. dev. of the noise in Eq. 1 was *σ* = 0.2.

### Scaling of the bias

Here we calculate the bias for small angles analytically and estimate how the bias scales with the model parameters. We use that in case of small Θ and the limit of small candidate angles *θ,* the mean square error can be Taylor expanded as:

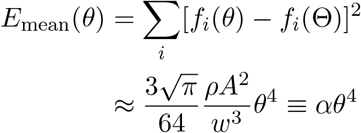

where we replaced the sum by an integral and where *ρ* is the coding density (the number of neurons per unit angle, *ρ* = *N*/2*π*). Similarly, the noise term on a given trial can expanded as

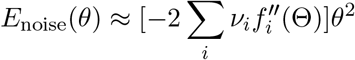

The coefficient in the square brackets is a Gaussian random variable with zero mean and a variance 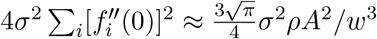. We are interested in the cases where the coefficient will be negative as these are the repulsive trials, which happens in half of the trials. The mean value of a Gaussian truncated below zero is 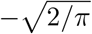 times the standard deviation, so that for these cases 〈(*E*_noise_ (*θ*)〉 ≈ − *βθ*^2^, with 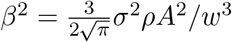.

The approximate location of the repulsed minimum is given by 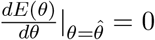, or 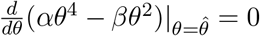, and thus 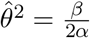. The bias in the other half of the trials is zero (purple traces in Fig. 2B), hence the total bias is 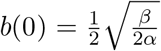. This yields the relation in the main text, Eq.3. The dependency of Eq.3 on all its parameters was confirmed numerically.

### Calculation of maximum likelihood estimate

Here we demonstrate how to calculate the distribution of estimates *P*(*θ̂*|Θ) of the ML estimator in a numerically exact manner. Given a noisy response r, we run over all candidate stimulus estimates and find the probability that it minimizes the Mean Square Error. Because the Mean Square Error is a smooth Gaussian process, and nearby E’s are correlated, we can finely discretize *θ*. We define a set of *M* candidate estimates (*θ*_1_,…,*θ_M_*). To calculate that the probability that a certain estimate *θ_m_* yields the lowest MSE, it is compared to the MSE that all other *M* − 1 estimates yield. We define the *M* − 1 dimensional set of MSE differences as **D***m* = *E*(*θm*) − *E*(Φ*m*), where Φ_*m*_ = {*θ*_1_,…, *θ_M_*}\*θ_m_*.

The distribution of differences **D**_*m*_ is a (*M* − 1)-dimensional multivariate normal distribution

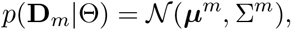

where *μ^m^* = *E*_mean_(*θm*) − *E*_mean_(Φ_*m*_) and the (*M* − 1) × (*M* − 1) covariance matrix has entries 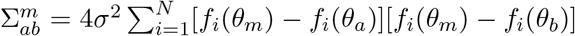. The probability that *θ_m_* has a lower MSE than all other candidate estimates, is

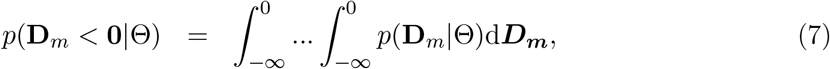

which is a multi-variate cumulative normal distribution.

While this orthant integral is not analytically tractable, efficient algorithms exist that calculate it to a high precision for values of *M* up to in the hundreds. We used the quasi-Monte Carlo integration function **mvnun** from Scipy (Genz, 1992, 1998) with *M* = 100 and *θ* = 0…*π* (using a larger *M* had negligible effects), and evaluated the integral for all values of *m.* This yields *P* (*θ̂*|Θ).

We note that while we applied it here to the estimation of Θ, this approach is general; it allows for arbitrary tuning curves and correlated Gaussian noise. It also extends to higher dimensional stimuli, but as one needs to discretize the stimulus space, a limitation is the efficient calculation of the integrals. Algorithms that calculate them for even higher dimensions exist (e.g.Azzimonti and Ginsbourger, 2016).

### Calculation of the Cramer-Rao bound

Here we show how the Fisher Information is calculated which we use to compare to the variance in the estimator. The Fisher Information matrix for additive, uncorrelated Gaussian noise is given by 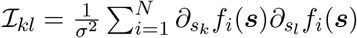. In the limit of dense tuning curves, the sum becomes an integral. While in the original s-coordinates the Information matrix has off-diagonal elements (Orhan and Ma, 2015), in the coordinates (Θ, *η*) it becomes diagonal

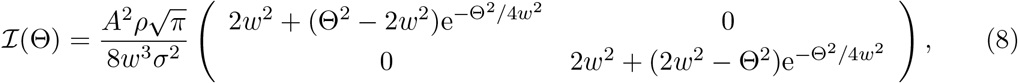

The diagonal nature confirms the intuition that the opening and the sum angles can be estimated independently. Further note that both information components depend on the opening angle Θ, but neither depends on the sum angle *η*. This is due to the rotation invariance of the problem w.r.t. *η*. Finally, the Fisher Information for Θ, that is 𝓘_11_, is zero for small Θ(Amari and Nakahara, 2005).

An estimator is called *efficient* if its decoding covariance satisfies the Cramér-Rao bound (CRB) (Rao, 1945; Cramér, 1946; Rao, 2008). In the oft studied case of un-biased, one dimensional estimators, the CRB is var(*θ̂*) ≥ l/𝓘 (Θ). In the case of biased vector parameters the CRB states that the matrix

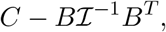

should be a positive definite matrix (Cover and Thomas, 1991; Kay, 1993). Here *C* is the covariance matrix of the stimuli that the estimator yields, *B* is the sum of the Jacobian matrix of **b** and the identity matrix. In our case this reduces to the bound in the main text.

### Fisher Information in max-coding model

For max-coding the Fisher Information is identical for both sum and difference angles, 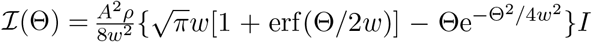, where *I* is the 2 × 2 identity matrix. This is a monotonic function in Θ. When there are two separate peaks in the population response (Θ≫*w*), the information is twice that when Θ= 0, where there is a single peak in the tuning.

## Supplementary figure

**Figure S1.**
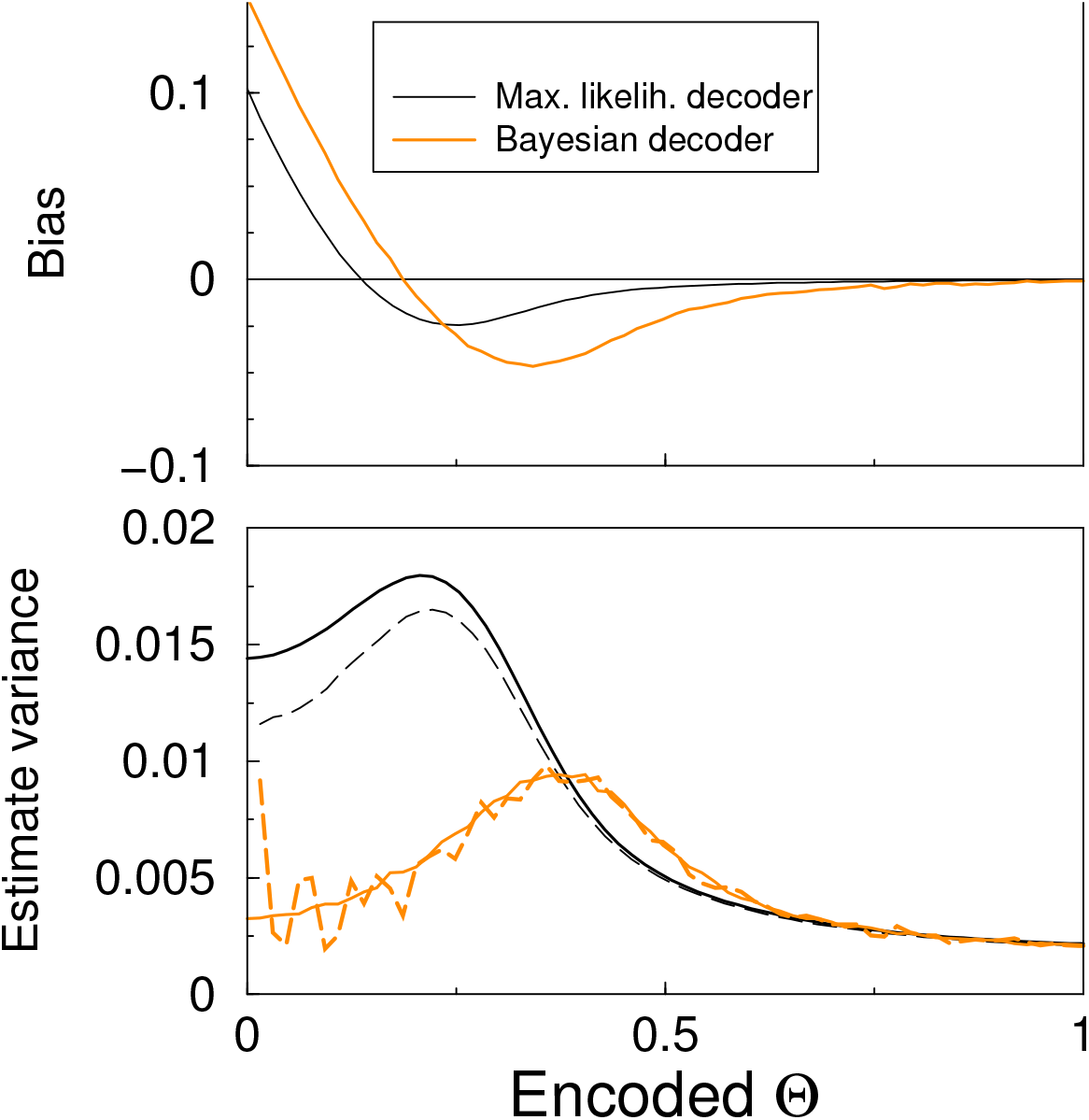
Bias and efficiency of a Bayesian decoder. Top: Bias in the estimate when using a Bayesian decoder (orange). Shown for comparison the ML decoder used in the main text (black). The biases are of comparable magnitude and share the biphasic character. Bottom: Standard deviation in the estimator. Dashed line shows the Cramer-Rao bound. The ML decoder is shown in black. Note that due to its dependence on the bias, the bounds for the estimators are different. The Cramer-Rao bound for the Bayesian decoder is more variable because it relies on a bias that needs to be extracted from simulations.

